# Differential Community Detection in Paired Biological Networks

**DOI:** 10.1101/147538

**Authors:** Raghvendra Mall, Ehsan Ullah, Khalid Kunjia, Halima Bensmail

## Abstract

**Motivation:** Biological networks unravel the inherent structure of molecular interactions which can lead to discovery of driver genes and meaningful pathways especially in cancer context. Often due to gene mutations, the gene expression undergoes changes and the corresponding gene regulatory network sustains some amount of localized re-wiring. The ability to identify significant changes in the interaction patterns caused by the progression of the disease can lead to the revelation of novel relevant signatures.

**Methods:** The task of identifying differential sub-networks in paired biological networks (A:control,B:case) can be re-phrased as one of finding dense communities in a single noisy differential topological (DT) graph constructed by taking absolute difference between the topological graphs of A and B. In this paper, we propose a fast two-stage approach, namely Differential Community Detection (DCD), to identify differential sub-networks as differential communities in a de-noised version of the DT graph. In the first stage, we iteratively re-order the nodes of the DT graph to determine approximate block diagonals present in the DT adjacency matrix using neighbourhood information of the nodes and Jaccard similarity. In the second stage, the ordered DT adjacency matrix is traversed along the diagonal to remove all the edges associated with a node, if that node has no immediate edges within a window. We then apply community detection methods on this de-noised DT graph to discover differential sub-networks as communities.

**Results:** Our proposed DCD approach can effectively locate differential sub-networks in several simulated paired random-geometric networks and various paired scale-free graphs with different power-law exponents. The DCD approach easily outperforms community detection methods applied on the original noisy DT graph and recent statistical techniques in simulation studies. We applied DCD method on two real datasets: a) Ovarian cancer dataset to discover differential DNA co-methylation sub-networks in patients and controls; b) Glioma cancer dataset to discover the difference between the regulatory networks of IDH-mutant and IDH-wild-type. We demonstrate the potential benefits of DCD for finding network-inferred bio-markers/pathways associated with a trait of interest.

**Conclusion:** The proposed DCD approach overcomes the limitations of previous statistical techniques and the issues associated with identifying differential sub-networks by use of community detection methods on the noisy DT graph. This is reflected in the superior performance of the DCD method with respect to various metrics like Precision, Accuracy, Kappa and Specificity. The code implementing proposed DCD method is available at https://sites.google.com/site/ raghvendramallmlresearcher/codes.

## 1 Background

In the modern era complex networks are ubiquitous. Their omnipresence is reflected in a myriad of domains including web graphs [6], road graphs [11], social networks [24, 42], financial networks [4] and biological networks [22, 27, 43]. Here we focus on biological networks but the caveats introduced in this paper apply to networks in other domains.

In network biology, particularly in cancer research, comparisons are performed on gene regulatory networks [57] and DNA co-methylation networks [56] obtained from the gene expression and DNA methylation profiles respectively of healthy and diseased tissues. The goal is to identify genes whose expression or methylation levels are significantly different between the conditions and can lead to discovery of novel molecular diagnostic and prognostic signatures. It was shown in [53, 1, 9] that the gene regulatory networks undergo some amount of localized re-wirings as cancer progresses.

One of the primary problems in cell biology is to infer regulatory networks, that capture the interactions between molecular entities from high-throughput data. An important challenge that needs to be addressed is how the cell changes its behaviour in response to changes in copy number or alterations such as driver somatic mutations or an external stimuli. The gene expression and methylation levels change due to the downstream effect of the de-regulation of the global behaviour of the cell in different conditions, for example different cancer subtypes [9]. Hence, it can be suggested that driver mutations regulate functional pathways described by different local re-wirings in the intrinsic gene regulatory networks.

The problem of detecting significant changes in paired biological networks is different from popular graph theory problems like graph isomorphism [46] and sub-graph matching [51] for which various graph matching and graph similarity algorithms [5, 30] exist and have been utilized in biological networks[55, 45]. This problem has primarily been addressed either in a statistical framework [37, 21, 50, 33] or from a community detection perspective [33, 10, 54, 23, 14, 32] in literature.

In statistics, a common statistic used to distinguish one graph from another is the Mean Absolute Difference (MAD), which is defined as: 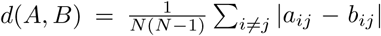 Here *a_ij_* and *b_ij_* are edge weights corresponding to the topological graphs of networks *A* and *B.* A topological graph captures first order interactions between the nodes in the network and can better apprehend subtle changes between two networks [49]. The MAD distance is equivalent to the Hamming distance [18] which has been widely used for comparing networks [7, 15]. The Quadratic Assignment Procedure (QAP) [37] defined as: 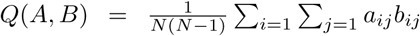 is another statistic used to identify association between networks. These statistics are often used in permutation-based procedures to detect significant difference between two networks. Ruan et al [50] showed that these statistics are not always sensitive to subtle topological variations and proposed a Generalized Hamming Distance (GHD) based statistic to measure the distance between paired biological graphs which outperforms MAD and QAP.

The GHD permutation distribution follows a normal distribution under the null hypothesis that networks *A* and *B* are independent for scale-free networks whose power-law exponent *α* should strictly satisfy: 1 ≤ *α* ≤ 2 or *α* ≥ 3. They also generated closed-form expression for p-values and devised a differential sub-network identification technique, namely dGHD, where they iteratively remove least different node. This is unlike previous differential network analysis techniques [15, 14, 17] and generate p-values by comparing the remaining subnetworks. Recently, a Closed-Form approach was proposed in [33] which is faster and more accurate than the dGHD technique for identifying statistically significant changes between paired networks as differential sub-networks. However, these statistical techniques are still computationally expensive and suffer from strict restrictions on the exponent of power-law for scale-free graphs. It was shown in [38] that biological networks are scale-free and usually have power-law exponents that satisfies: 0 *< α* ≤ 2 which is not always within the restrictions acceptable for dGHD and Closed-Form techniques.

The problem of community detection in graphs has received wide attention from several perspectives [16, 3, 48, 47, 44, 36, 34, 35, 29] and have also been applied to biological networks. Methods such as jActiveModules [10] and the Spinglass algorithm [47] have been applied to discover biologically meaningful modules such as protein complexes, disease associated clusters of genes, etc. as shown in [54, 23]. The problem of identifying differential sub-networks in paired biological networks can be re-formulated as one of finding heavy sub-networks, or dense modules, on a single differential topological (DT) graph obtained by taking the absolute difference in the edge weights between the topological graph of network A and the topological graph of network B i.e. DT(*A*, *B*)*_ij_* = *|a_ij_* − *b_ij_*|, ∀*i*, *j* ∊ *V.* This problem is equivalent to identifying communities in the DT graph. The notion of communities mean that nodes within one community are densely connected to each other and sparsely connected to nodes outside that community. Large-scale networks consist of several such communities. Hence, community detection is equivalent to finding dense block diagonals in the DT adjacency matrix. However, the DT graph can suffer from noise caused by interactions between nodes which are not part of differential sub-networks (referred further as *non-differential nodes*) and nodes which are part of differential sub-networks (referred further as *differential nodes*) which are just one hop away in either network A or B but not in both. This leads to spurious connections around the block diag onals present in the DT adjacency matrix. Community detection techniques like Louvain [3], Infomap [48] and Spectral [34] method can be applied to the obtain communities/modules with differential nodes with having perfect recall but suffer from very low precision due to false recognition of non-differential nodes as part of differential sub-networks.

The problem of identifying communities in the DT graph such that the nodes comprising the communities are part of differential sub-networks between paired biological networks (*A, B*) is unlike the traditional module based differential network analysis as shown in [14, 32]. In traditional module based differential network analysis, modules are detected at first in weighted gene co-expression networks (WGCNA) [14] obtained from gene expression data for case and controls. The modules are then compared using either Jaccard co-efficient (MOda) [32] or additional genetic marker data (WGCNA) [14] is utilized to differentiate the modules. The advantage of these methods is that by focusing on modules rather than on individual gene expressions, they can greatly alleviate the multiple-testing problem inherent in micro-array data analysis. However, our goal is to identify the difference between the paired biological networks as dense modules/communities rather than comparing the modules in the paired biological networks. For example, say minor localized changes within two modules in the original biological networks together form a differential sub-network. The method proposed in this paper will be able to identify these changes as a differential community which might otherwise not be detected by WGCNA or MOda.

In this paper, we propose a novel two-stage approach, namely Differential Community Detection (DCD), to identify differential sub-networks in paired biological networks as communities from the original nosiy DT graph. The proposed DCD method overcomes the restrictions on power-law exponents for scale-free graphs implied by statistical techniques and retains the advantage of greatly reducing the burden of multiple-testing from module based differential network analysis techniques. We applied our DCD method on two real datasets, an ovarian cancer dataset to discover differential DNA co-methylation sub-networks in patients and controls, and a glioma cancer dataset to discover the difference between the regulatory networks of IDH-mutant and IDH-wild-type.

## 2 Method

The proposed DCD approach consists of two primary stages: In the first stage of DCD, the proposed method re-orders the nodes of the DT graph to generate approximate block diagonals inherently present in the DT adjacency matrix. It utilizes the neighbourhood information from the DT graph for all the nodes and a notion of similarity based on the Jaccard index [31]. In the second stage of DCD, the ordered yet noisy DT adjacency matrix is traversed along the diagonal to remove all the edges associated with a node, if that node has no immediate edges within a window. This is because the ordered DT adjacency matrix is already comprised of block diagonals and nodes which are not part of block diagonals are the ones causing spurious connections in the DT graph. We then pick out such nodes and remove all the edges associated with these nodes. Finally, we apply community detection techniques like Louvain [3], Infomap [48] and Spectral [34] methods on this de-noised DT graph to discover the differential subnetworks as communities. Figure 1 illustrates all the steps involved in the DCD algorithm and its comparison with direct application of community detection techniques on noisy DT graph to locate differential sub-networks on a pair of simulated random-geometric (RG) networks.

**Figure 1:**
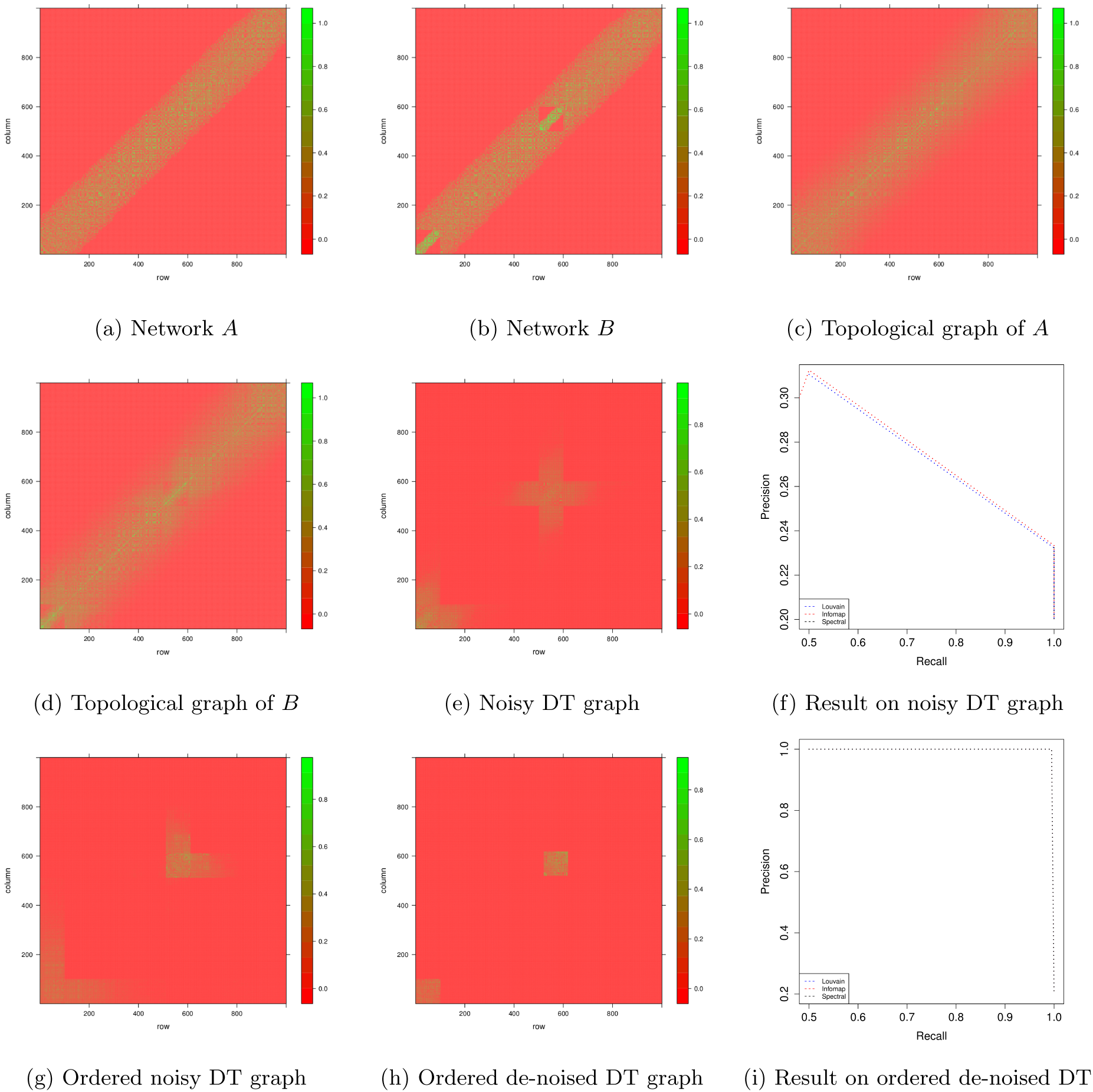
Illustration of DCD method and its benefit over directly using community detection methods on noisy DT graph. Figure 1a represents a random-geometric network *A* with 1, 000 nodes and Figure 1b represents another random-geometric network *B* where the nodes 1 to 100 and nodes 500 to 600 have different interaction pattern from network *A.* Figures 1c and 1d correspond to the topological graphs of network *A* and *B.* Figure 1e shows the noisy differential topological (DT) graph obtained from topological graphs of *A* and *B.* Figure 1f evaluates the result of 3 state-of-the-art community detection techniques on the noisy DT graph to detect differential sub-networks w.r.t. precision and recall metrics. Figure 1g illustrates the ordered noisy DT graph obtained from first stage of DCD approach. Figure 1h demonstrates the de-noised DT graph generated after the second stage of DCD method. Figure 1i showcases the efficiency of 3 different community detection methods to identify the differential sub-networks from the de-noised DT graph.

### 2.1 Ordering the Noisy DT graph

The goal of first stage of DCD method is to detect sets of nodes which have higher similarity with each other in comparison to other nodes by following an iterative procedure to order the nodes in the adjacency matrix of the original DT graph *G*(*V, E*). The total number of nodes in the DT graph is represented as *N* = *|V|.* This iterative process is essential as nodes are not usually ordered in the *G*(*V, E*) and the inherent block diagonals have to be discovered. It is important to locate approximate block diagonals as it is a necessary condition for the second stage of DCD approach. We define *d*(*v_i_,V^t^*) as degree of the node *v_i_* ∊ *V^t^*, where *V^t^* represents the set of nodes to be investigated at iteration *t.* During the first iteration, we identify the node with highest degree i.e. 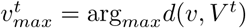 using the topology of *G*(*V, E*) and calculate its Jaccard similarity w.r.t. all the nodes in DT graph. Mathematically, it is defined as:

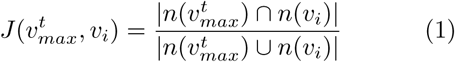

Here 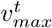 is the node with highest degree during iteration *t, v_i_* ∊ *V, n*(*·*) represents the immediate neighbourhood set of a node and |·| represents the cardinality function. The Jaccard co-efficient of all the nodes that don’t share a specified number of neighbours (*θ*) with 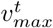 is set to 0. This threshold *θ* is a tunable parameter representing the minimum size of a block diagonal to be considered as a differential community in the DT graph. We then sort all the nodes having non-zero Jaccard similarity with 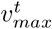 in decreasing order and break ties based on degree where higher degree nodes are placed closer to 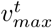. These ordered nodes and their corresponding edges results in the first approximate block diagonal *ABD^t^* which is preserved in ODT, representing the adjacency matrix of ordered noisy DT graph. *ABD^t^* is an approximate block diagonal because nodes with spurious connections are still present and associated with *ABD^t^* as highlighted in Figure 1g.

During further iterations (*t >* 1), an additional step is performed to re-order the nodes which are common between the *ABD^t^*^−1^ and *ABD^t^.* The order of common nodes whose Jaccard similarity was higher with the previous 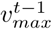 are unchanged and these nodes are removed from *ABD^t^.* However, nodes which are common with *ABD^t^*^−1^ but have higher Jaccard similarity with 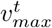 are removed from *ABD^t^*^−1^ while their order is retained in *ABD^t^.* This iterative process is greedy by nature, as in any iteration *t* we compare only *ABD^t^*^−1^ with *ABD^t^*, and stop when either all the nodes in the *G*(*V, E*) are part of some approximate block-diagonal or degree of 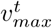 is 0, which means we are left with only isolated nodes in the *G*(*V, E*). Algorithm 1 summarizes this procedure.

#### Algorithm 1: Ordering Noisy DT graph

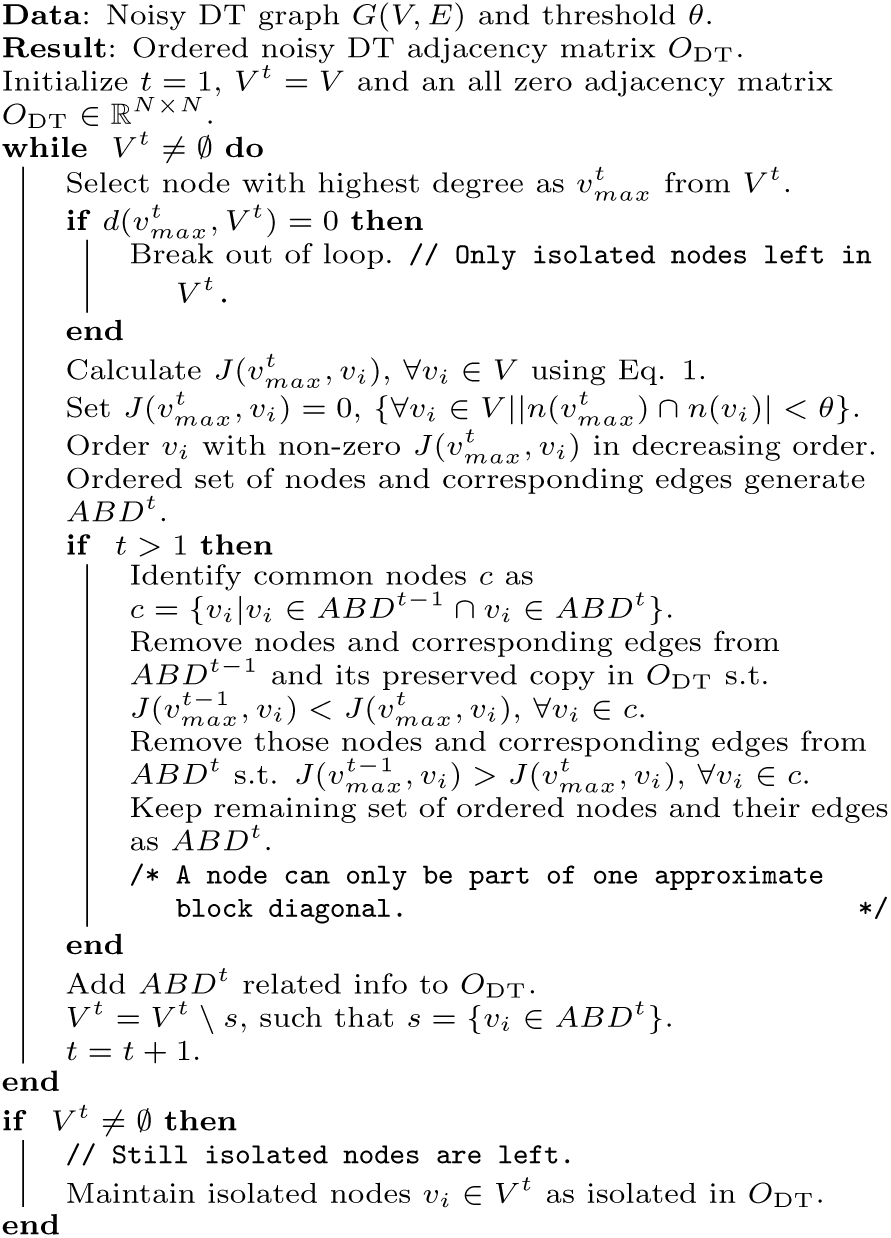

### 2.2 De-noising the DT graph

Once we have obtained *O*_DT_ as shown in Figure 1g, we prune out spurious edges associated with nodes which are falsely recognized as part of block diagonals in the previous step. We traverse the landscape of the *O*_DT_ matrix, for example in Figure 1g from left to right and bottom to up, along the diagonal. Since we have already identified approximate block diagonals (*ABD’s*) in *O*_DT_, our premise is that if we traverse along the diagonal and pick a node *v_i_* at random, there should be some immediate edges within *θ* to the left and to the right (below and above due to symmetry) in the landscape of *O*_DT_ for it to be a differential node in *ABD.* This means that *d*(*v_i_, V_i_*_−*θ*_) and *d*(*v_i_, V_i_*_+*θ*_) have to be non-zero at the same time. Here *V_i_*_−*θ*_ and *V_i_*_+*θ*_ represent the neighbourhood up to *θ* nodes to the left and right of *v_i_.* A non-differential node can be part of *ABD* due to spurious connections with the differential set of nodes present in *ABD.* We then remove all the edges associated with such nodes from *O*_DT_ to generate the de-noised ordered DT graph i.e. *D*_DT_. The proposed process leads to de-noised block diagonals *BD* in *D*_DT_ instead of having *ABD* as shown in Figure 1h. Algorithm 2 summarizes the de-noising procedure.

#### Algorithm 2: De-noising the DT graph

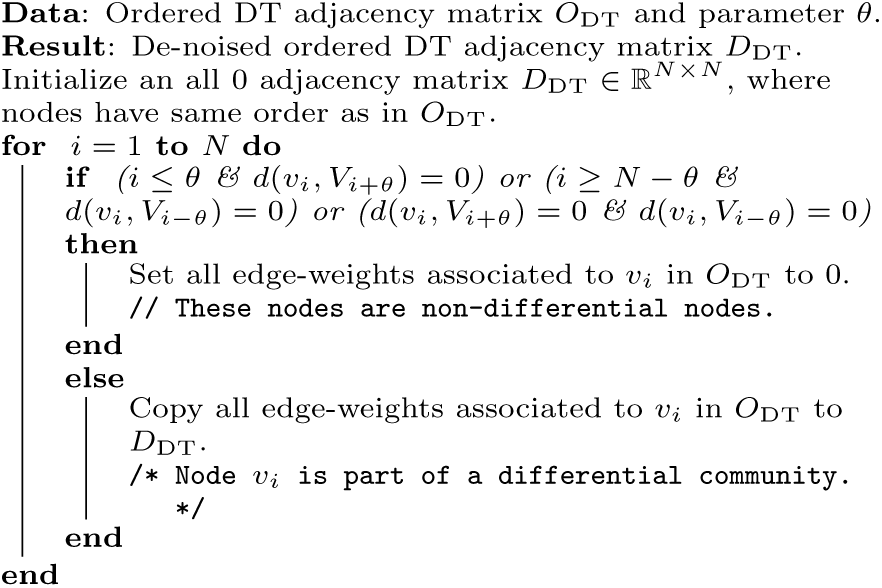

We can now run state-of-the-art community detection algorithms [34, 3, 48] to distinguish the *BD*’s in *D*_DT_ as differential communities in paired biological networks. The overall time complexity of proposed steps is O(*tN* log *N* + *tEd_μ_*), where *t* is number of iterations in Algorithm 1, *E* represents number of edges and *d_μ_* represents the average degree of a node in DT graph. Algorithm 3 provides an overview of the proposed DCD approach.

## 3 Simulated Experiments & Results

We performed multiple simulated experiments on paired random-geometric (RG) and paired scale-free networks under different experimental settings. All the experiments were repeated 10 times for each experimental setting.

#### Algorithm 3: Differential Community Detection (DCD) approach for paired biological networks

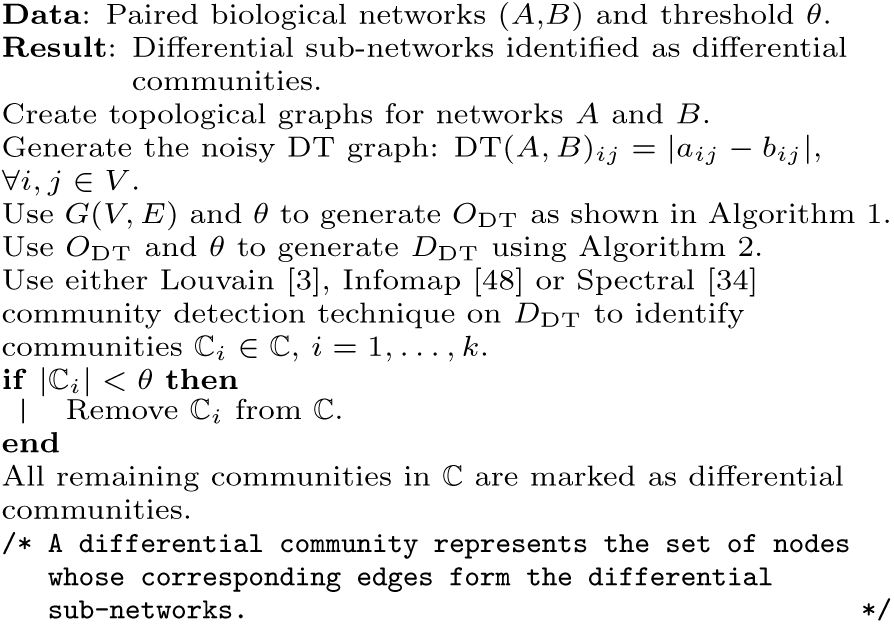

In an RG network nodes are generated by uniformly sampling *N* points on [0,1]^2^. An edge is drawn between points if the euclidean distance between the points is less than a parameter *v.* This parameter *v* controls the density of the RG network where smaller values of *v* result in sparse networks while larger values of *v* result in dense networks. We performed two set of experiments on RG networks. In the first case, we generated RG network *A*_1_ with *N* = 1, 000 and *v* = 0.15. Network *B* is obtained by permuting first 100 nodes in network *A.* Thus, these first 100 nodes form the differential sub-network for the paired RG networks *A*_1_ and *B*_1_.

In the second case, we again used *N* = 1, 000 and *v* = 0.15 to generate network *A*_2_. We then create a small dense RG network with 100 nodes using 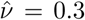. Network *B*_2_ was generated by replacing first 100 nodes in network *A*_2_ with the small dense sub-network. These 100 nodes form the differential sub-network for the paired networks *A*_2_ and *B*_2_. Such a mechanism can appear in real-life networks, for example, in case of cancer the transcription activity of some set of genes might get enhanced or suppressed generating more or fewer edges in a sub-network of the gene or DNA methylation network. We performed similar set of experiments using density parameter *v* = 0.3 and permuting first 100 nodes, using density parameter *v* = 0.3 and adding more edges to first 100 nodes using revised density parameter 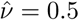 on paired RG networks.

We also conducted experiments on undirected scale-free graphs, hereby referred as Power-Law (PL) networks, using *N* = 1000 and *E* = 10, 000 with varying power-law exponents *α* = {1, 1.5, 2} respectively. We permuted the first 100 nodes of each PL network (*A*) to form the permuted network (*B*). The proposed DCD method has one tunable parameter *θ.* In Figure 2, we illustrate the effect of *θ* on the area under the precision-recall curve. From Figures 2a, 2b, 2e, 2f, 2i and 2j, we can observe that for smaller values of *θ* ({3, 5}), the area under precision-recall curves are relatively lower in comparison to those for higher values of *θ.* This is due to the fact that for smaller values of *θ*, we are allowing smaller sized communities to be distinguished as differential sub-networks. This can force to break the natural block diagonals inherently present in the DT graph and reduce the number of true positives (i.e. nodes which are actually part of differential sub-networks) leading to lower precision and recall. At the same time, smaller values of *θ* allow non-differential nodes with few spurious connections to differential nodes to be falsely identified as part of differential sub-networks resulting in lower precision. For higher values of *θ* ({7,9}), the area under precision-recall curves shows less variance and converges to nearly perfect result (≈ 1) as depicted in Figures 2c, 2g,2h, 2k and 2l.

**Figure 2:**
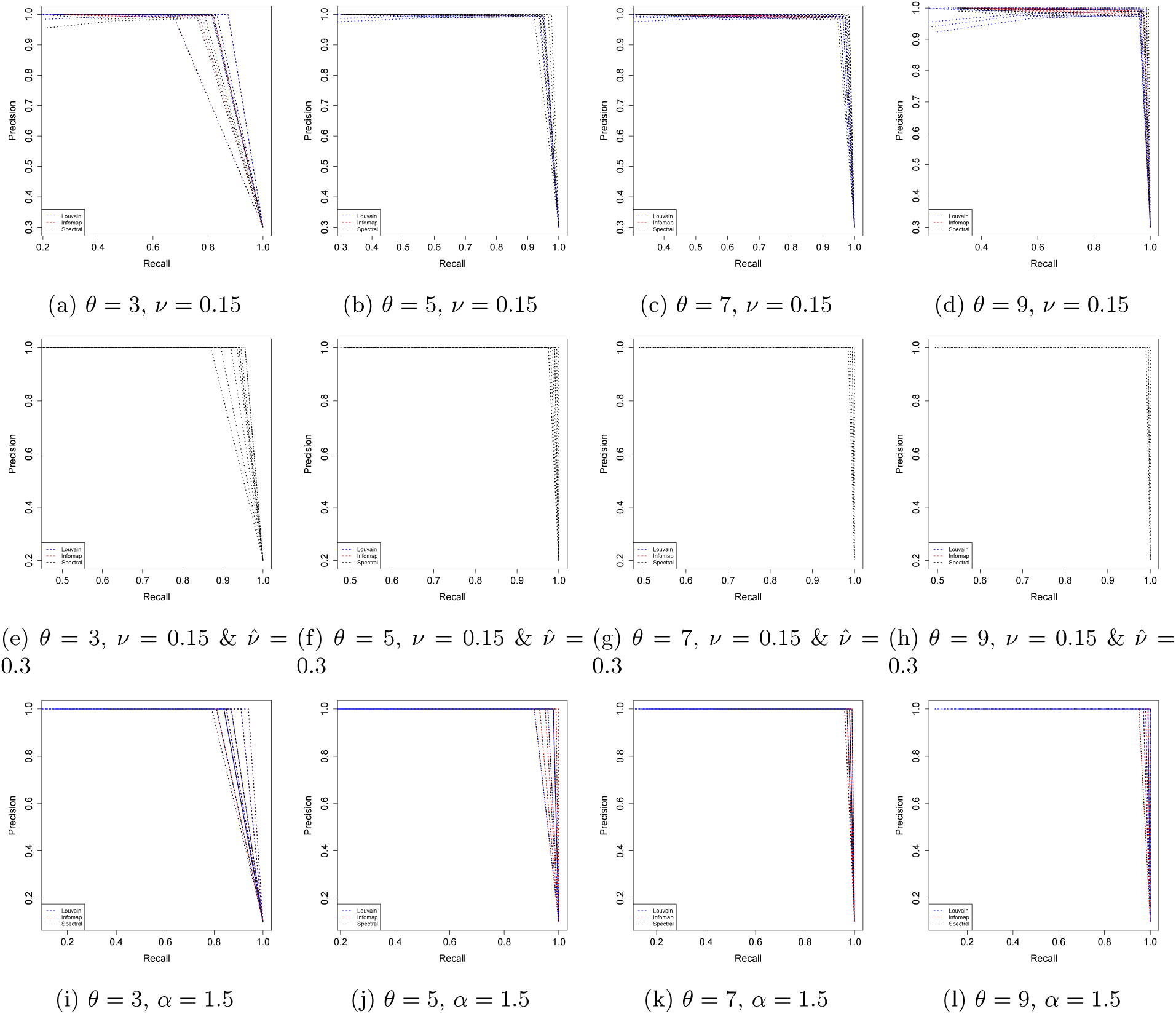
Area under the precision-recall curves for different values of threshold *θ* for various experimental settings. We demonstrate the area under precision-recall curves using the proposed steps of DCD approach with either Louvain or Infomap or Spectral community detection method. Figures 2a,2b,2c and 2d show the role of parameter *θ* on precision-recall values for paired RG networks (*ν* = 0.15) where first 100 nodes are permuted. Figures 2e, 2f, 2g and 2h illustrate how the area under precision-recall curves vary with threshold *θ* for paired RG networks (*ν* = 0.15) where the sub-network corresponding to first 100 nodes have higher density 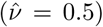. Similarly, Figures 2i, 2j, 2k and 2l describes the role of variable *θ* on precision-recall values for paired PL networks (*α* = 1.5) where the first 100 nodes are permuted.

Table 1 encapsulates a comprehensive comparison of the proposed DCD approach, where the community detection technique used in DCD is either Louvain [3] or Infomap [48] or Spectral [34], with statistical techniques like dGHD [50] and Closed-Form [33] approach and direct application of community detection methods like Louvain, Infomap and Spectral on the noisy DT graph to detect differential subnetworks in the simulated experiments. We used the threshold *θ* = 7 in the DCD approach for all comparisons as the area under precision-recall curves shows less variance and converges to nearly perfect value (≈ 1) in all the simulated experimental settings for this threshold as depicted in Figure 2. For nearly all PL graph experiments, if we directly apply community detection methods on the noisy DT graph, they identify all the nodes in the network as part of differential sub-network as depicted from evaluation metrics in Table 1.

**Table 1:**
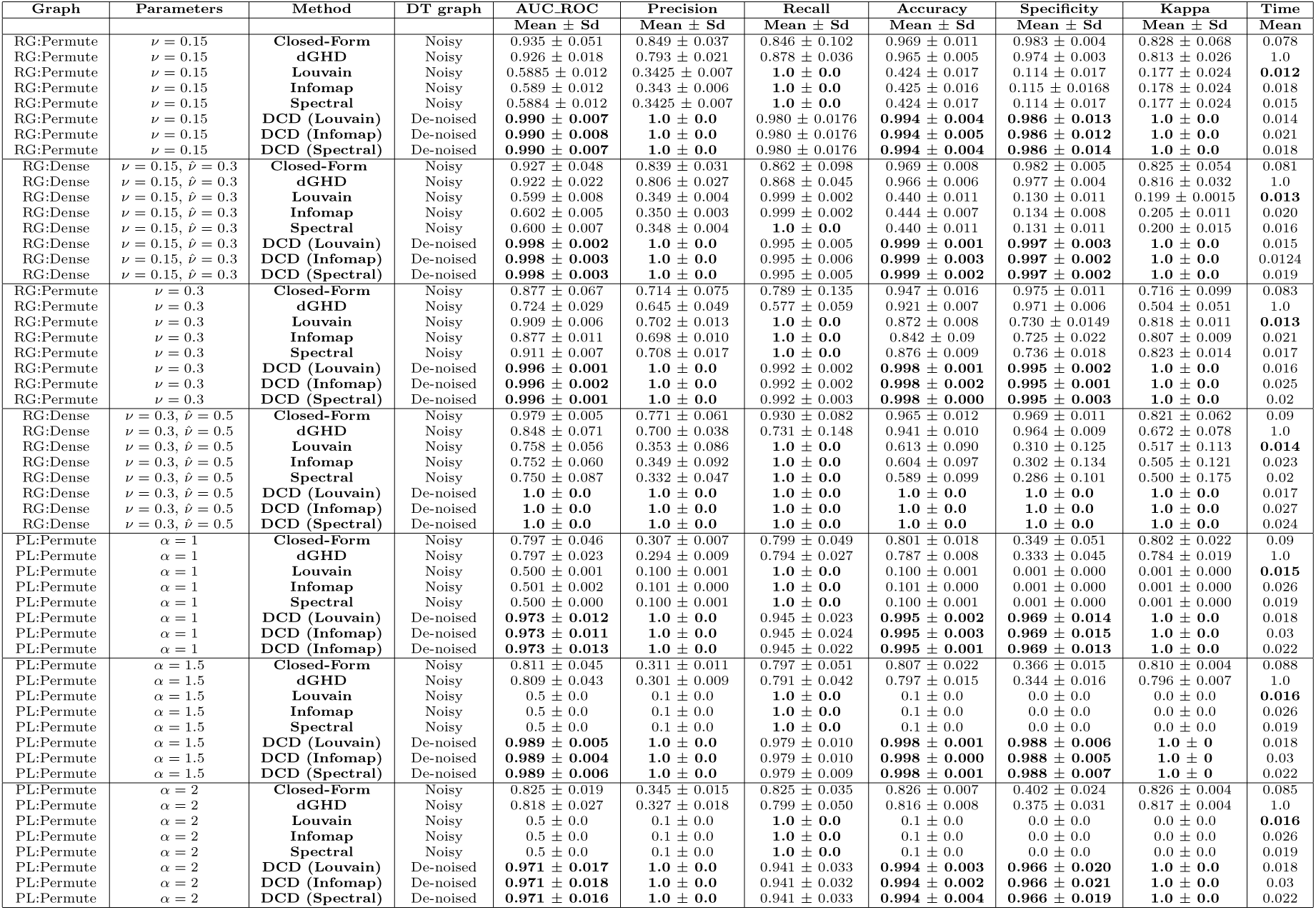
Comparison of proposed DCD approach with Closed-Form [33] and dGHD [50] statistical techniques and direct application of community detection methods like Louvain [3], Infomap [48] and Spectral [34] on nosiy DT graph to identify differential sub-networks in paired simulated networks for various settings. Here RG:Permute represents RG networks where first 100 nodes are permuted and form differential subnetwork. Similarly, PL:Permute is used for experiments on PL graphs where first 100 nodes are permuted and constitute the differential sub-network. RG:Dense depicts RG networks, where first 100 nodes have higher density in network B in comparison to network A and make-up the differential sub-network. Time is represented as fraction w.r.t. the computational time of most expensive method (dGHD). Best results are highlighted in bold. The proposed DCD approach can robustly identify differential sub-networks in all simulated experimental settings. It performs the best for evaluation metrics: AUC_ROC (area under ROC curve), Precision, Accuracy, Kappa and Specificity.

## 4 Application to comethylation networks in ovarian cancer

We applied our proposed DCD approach, with parameter *θ* set to 7, on co-methylation networks generated from ovarian cancer dataset [52]. Thus, the smallest community in DT graph should comprise at least 7 nodes. The ovarian cancer dataset consists of methylation profiles for 27, 578 CpG islands of 540 women, of which 266 cases were from postmenopausal women with ovarian cancer and 274 were healthy controls with similar age as that of cases. In our analysis, we have compared case and control DNA co-methylation networks to identify differential sub-networks.

The pre-processed dataset was downloaded from GEO (repository number GSE19711). The original data was collected using Illumina Infinium 27k Human DNA methylation Beadchip v1.2. Since there were no missing or negative values for the intensity of the methylated (*M*) and unmethylated (*U*) alleles, beta values corresponding to each CpG probe were computed as: 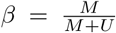 as in [50]. We followed the quality control procedure as originally introduced in [52]. Then principal component analysis (PCA) was applied to the beta values for detection and removal of outliers. After quality control, 243 case samples and 214 control samples remained for further analysis. Networks for case and control samples were created by treating each probe as a node. Edges between the nodes represent strong correlation and were inferred following [19]. Adjacency measure Ω*_ij_* was computed for each pair of nodes (*i* and *j*) as 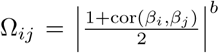, where cor(*β_i_*, *β_j_*) represents Pearson’s correlation coefficient between beta values observed at *i*^th^ and *j*^th^ CpG sites. The exponent *b* was set to 12 to emphasize more on higher positive correlations [57]. An edge exists if Ω*_ij_* value was higher than 0.2. The resulting control network has 73, 145 edges and case network has 102, 799 edges. Each of these networks follows a scale-free network model as shown in Figure 3.

**Figure 3:**
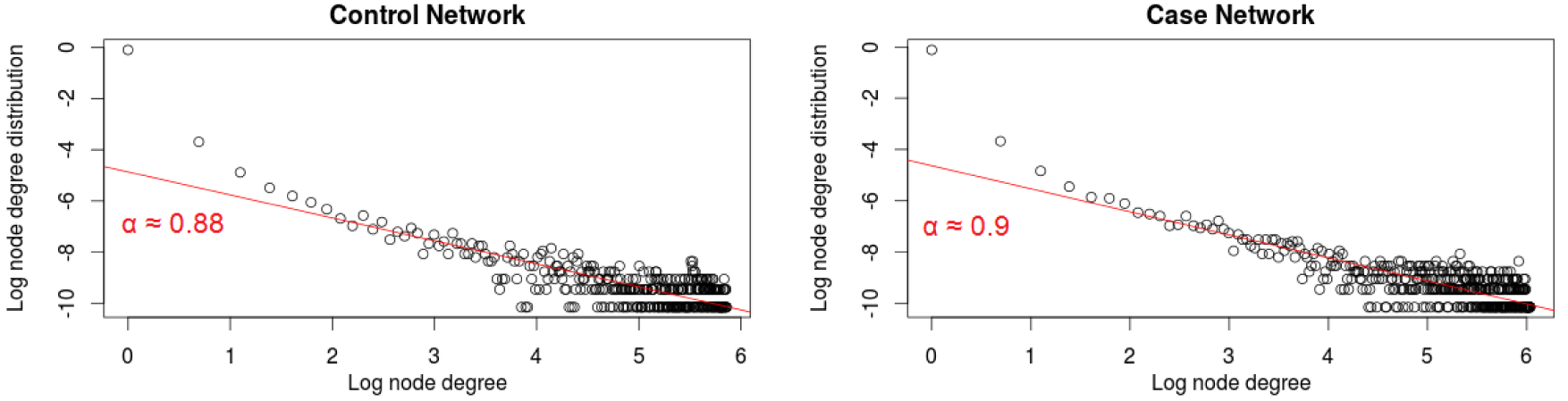
Degree distribution of nodes for control and case co-methylation networks. Since *α <* 1 for both the networks, state-of-the-art statistical techniques cannot be applied on these paired networks for differential sub-network analysis.

Our approach detected differential sub-networks comprising of a total of 1, 893 nodes. We used Louvain [3] method for detection of communities in the differential case and control sub-networks. Nine communities were detected in the case differential subnetwork out of which seven are also present in the control differential sub-network as shown in Figure 4.

**Figure 4:**
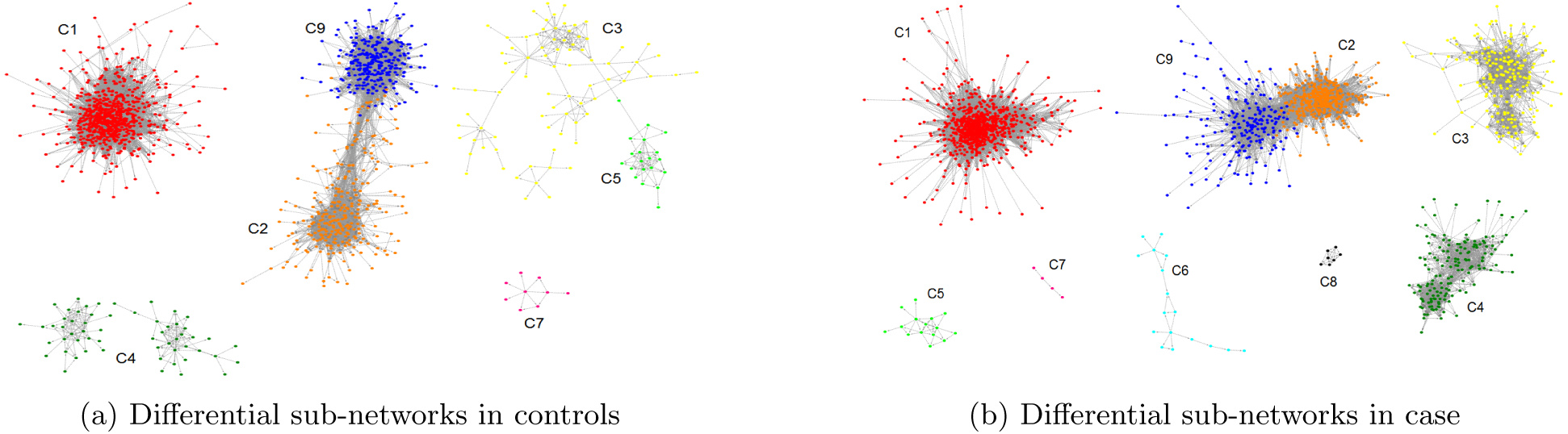
DNA co-methylation differential sub-networks. Cluster C7 is a special case. Even though it comprises of less than 7 nodes in the case sub-network, it consists of 9 nodes in control sub-network and has very different topography in the two sub-networks. As a result, it appears as a differential community of size greater than 7 in the de-noised DT graph. Clusters C6 and C8 are not present in the control sub-network.

We investigated the biological meaning of the sub-networks by identifying enriched Gene Ontology (GO) terms. We used R package GOstats [13] to identify Biological Processes (BP) and Molecular Functions (MF). The hypergeometric test detected 711 BP and 100 MP statistically significant terms enriched in the sub-networks at 5% significance level. The top three BPs were regulation of myeloid cell apoptotic process, myeloid cell apoptotic process, and establishment of protein localization to organelle. The top three MFs were protein binding, peroxidase activity and glycosaminoglycan binding. Furthermore, we identified 16 significantly enriched KEGG pathways at 5% significance level including transcriptional mis-regulation in cancer, hematopoietic cell lineage, and pathways in cancer using DAVID [20].

We detected probes with significant changes in mean methylation levels using the t-test. We found 5,098 significantly differentially methylated CpGs at 5% significance level after FDR correction for multiple testing [2]. Table 2 summarizes the number of probes, differentially methylated probes (*q_i_*), density ratio between control and case sub-networks (*R_i_*), and distribution of enriched GO terms and KEGG pathways in the identified communities.

**Table 2:**
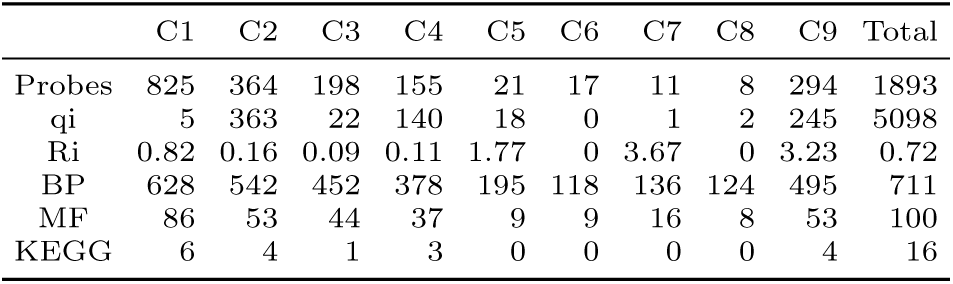
DNA co-methylation networks: a summary of different communities detected by DCD approach.

## 5 Application in Glioma Cancer

We also applied the DCD approach, with parameter *θ* set to 7, on gene regulatory networks (GRN) generated from the TCGA pan-glioma dataset [33]. The TCGA pan-glioma dataset includes 1, 250 samples (463 IDH-mutant and 653 IDH-wild-type), 583 of which were profiled with Agilent microarray and 667 with RNA-Seq Illumina HiSeq (REF) downloaded from the TCGA portal. The batch effects between the two platforms were corrected using the COMBAT algorithm [25]. The final gene expression data includes 12,985 genes and 1, 250 samples. From this data, we inferred the GRN for the two different glioma sub-types using the ARACNe [39] algorithm as in [33]. In our analysis, we compared the GRNs of IDH-mutant and IDH-wild-type to identify sub-networks of transcription factors (TFs) having a different regulatory program in these two major conditions.

The ARACNe networks were intersected with an active binding network based on the presence of binding sites in the promoter of a target gene. The active binding network is reconstructed for 2,532 unique motifs corresponding to 1,203 unique TFs [26, 40, 28]. A binding relationship is considered active if the TF motif signal is significantly (FDR *<* 0.05) over-represented in the target promoter region (∓ 5kbp TSS, hg19) and, in the same position (at least 1bp overlapping), chromatin state is classified as open by Hidden Markov Model proposed in [12]. The active binding network consists of 6,652, 518 overlapping active sites resulting in 1,959,125 unique TF associations between 1, 203 TFs and 51, 705 targets.

The final pruned networks are then obtained by considering the common sub-network of active binding and functional ARACNE networks. They consists of 13,683 unique connections for IDH-mutant and 14,158 for IDH-wild-type between TF-TF and TF-target. The number of TFs was reduced to 457 when intersected with the 12, 895 genes of our combined expression matrix. We then apply the proposed DCD approach on the noisy DT graph *G*(*V, E*) obtained by taking the absolute difference between the topological graphs of IDH-mutant and IDH-wild-type. The DCD technique discovered a total of 262 TFs as part of 7 differential communities using the Louvain [3] method in *G*(*V, E*).

We further investigated these communities by considering the regulons of all the TFs associated with each such community Ci in the corresponding IDH-mutant and IDH-wild-type GRN. The regulon of a TF is defined as its neighbourhood in the GRN. We probed the regulons of all TFs present in a community to detect enriched GO terms using DAVID [25]. We found 15 and 17 statistically significant biological processes (BP) at a 5% significance level using the regulons of TFs in *C*_1_ for IDH-mutant and IDHwild-type GRNs respectively. We also located 50, 14, 9, 21, 51 and 40 significant BPs for *C*_2_, *C*_3_, *C*_4_, *C*_5_, *C*_6_ and *C*_7_ respectively in IDH-mutant GRN. Similarly, we unearthed 71, 11, 4, 20, 48 and 20 significant BPs for *C*_2_, *C*_3_, *C*_4_, *C*_5_, *C*_6_ and *C*_7_ respectively in IDH-wildtype GRN.

We utilized the output from DAVID for each *C_i_* in the IDH-mutant and IDH-wild-type GRN as input to Enrichment Map tool [41] in Cytoscape. This tool provides a visualization for functional enrichment associated with BPs in *C_i_* and allows comparison between enrichment results for two different conditions (IDH-mutant and IDH-wild-type). Figure 5a illustrates the difference between the enrichment results of *C*_1_ in IDH-mutant and IDH-wild-type case. Similarly, Figure 5b compares the enrichment results of *C*_3_ in IDH-mutant and IDH-wild-type.

**Figure 5:**
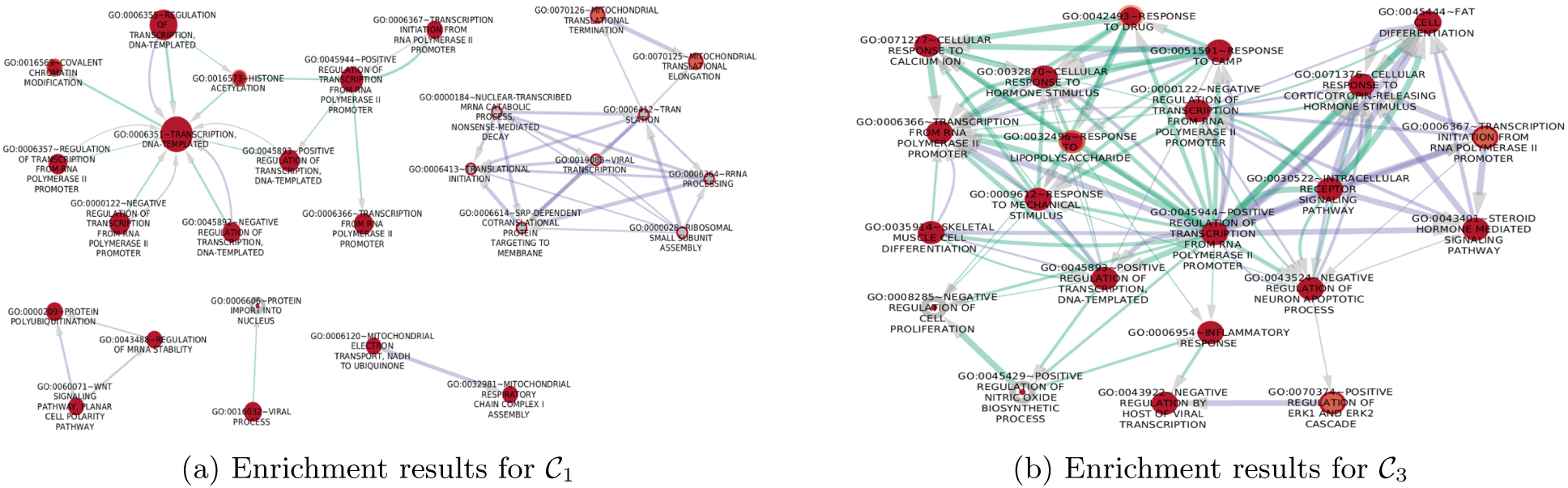
Comparison of enrichment results of IDH-mutant and IDH-wildtype for differential communities *C*_1_ and *C*_3_. Here the nodes correspond to the BPs and red circle size is proportional to number of genes in IDH-mutant associated with that BP. Similarly, the grey circle size in a node (BP) corresponds to the number of genes in IDH-wild-type related to that BP. Edge size corresponds to the number of genes that overlap between the two connected BPs. Green edges correspond to IDH-mutant while purple edges represent interaction between BPs in IDH-wild-type.

Interestingly, the differential community *C*_1_ is enriched with functions related to epigenetic changes such as Chromatin Modification and Histone Acetylation. Ceccarelli et al showed in [8] that the main difference between IDH-mutant and IDH-wild-type gliomas is the characteristic hyper-methylation phenotype (G-CIMP) which has a favourable prognosis both in high grade and low grade gliomas. Conversely, the *C*_3_ reveals enrichments which are specific of IDH-wild-type gliomas such as proliferation and activation of inflammatory response. Therefore, the DCD approach is not only able to identify known but also potential novel enrichments which need to be investigated further, in the two pathological conditions. Additional supplementary information is provided at https://sites.google.com/site/raghvendramallmlresearcher/codes.

## 6 Conclusion

We propose a fast two-stage DCD approach to identify differential sub-networks in paired biological graphs. The proposed method performs node ordering using neighbourhood information of nodes and Jaccard similarity to detect approximate block-diagonals. It de-noises the ordered noisy differential topological graph by traversing its landscape along the diagonal. Finally, differential sub-networks are identified using community detection algorithms. We showcased the effectiveness of proposed approach w.r.t. various statistical techniques and direct application of community detection methods for a myriad experimental settings using evaluation metrics like Precision, Accuracy, Kappa and Specificity. The DCD approach identified several meaningful biological processes and molecular functions on ovarian cancer dataset. Similarly, using DCD, we singled out some functional pathways that are different between the IDH-mutant and IDH-wild-type subtypes in case of glioma cancer.

## References

[1] Ahern, T., Horvath-Puho, E., Spindler, K., Sorensen, H., Ording, A., and Erich-sen, R. Colorectal cancer, comorbidity, and risk of venous thromboembolism: assessment of biological interactions in a Danish nationwide cohort. British Journal of Cancer 114, 1 (2016), 96–102.

[2] Benjamini, Y., and Yekutieli, D. The control of false discovery rate in multiple testing under dependency. Annals of Statistics 29 (2001), 1165–1188.

[3] Blondel, V. D., Guillaume, J.-L., Lambiotte, R., and Lefebvre, E. Fast unfolding of communities in large networks. Journal of statistical mechanics: theory and experiment 2008, 10 (2008), P10008.

[4] Boginski, V., Butenko, S., and Pardolas, P. M. Statistical analysis of financial networks. Computational Statistics and Data Analysis 48, 2 (2005), 431–443.

[5] Brandes, U., and Eriebach, T. Network Analysis: Methodological Foundations. Springer 3418 (2005).

[6] Broder, A., Kumar, R., Maghoul, F., Raghavan, P., Rajagopalan, S., Stata, R., Tomkins, A., and Wiener, J. Graph structure in the web. Comput. Netw. 33, 1-6 (2000), 309–320.

[7] Butts, C., and Carley, K. Canonical labeling to facilitate graph comparison. Tech. rep., Carniege Mellon University, 1998.

[8] Ceccarelli, M., Barthel, F. P., Malta, T. M., Sabedot, T. S., Salama, S. R., Murray, B. A., Morozova, O., Newton, Y., Radenbaugh, A., Pagnotta, S. M., et al. Molecular Profiling Reveals Biologically Discrete Subsets and Pathways of Progression in Diffuse Glioma. Cell 164, 3 (Feb. 2016), 550–563.

[9] Ceccarelli, M., Cerulo, L., and Santore, A. De novo reconstruction of gene regulatory networks from time series data, an approach based on formal methods. Methods 69, 3 (Oct 2014), 298–305.

[10] Dittrich, M. T., Klau, G. W., Rosenwald, A., Dandekar, T., and Müller, T. Identifying functional modules in protein–protein interaction networks: an integrated exact approach. Bioinformatics 24, 13 (2008), i223–i231.

[11] Erath, A., Lchl, M., and Axhausen, K. Graph-theoretical analysis of the swiss road and railway networks over time. Networks and Spatial Economics 9, 3 (2009), 379400.

[12] Ernst, J., and Kellis, M. Chromhmm: automating chromatin-state discovery and characterization. Nature methods 9, 3 (2012), 215–216.

[13] Falcon, S., and Gentleman, R. Using GOstats to test gene lists for GO term association. Bioinformatics 23, 2 (2007), 257–258.

[14] Fuller, T., Ghazalpour, A., Aten, J., Drake, T., Lusis, A., and Horvath, S. Weighted Gene Co-expression Network Analysis Strategies Applied to Mouse Weight. Mammilian Genome 18, 6 (2007), 463–472.

[15] Gill, R., Datta, S., and Datta, S. A statistical framework for differential network analysis from microarrya data. BMC: Bioinformatics 11, 1 (2010), 95.

[16] Girvan, M., and Newman, M. E. Community structure in social and biological networks. Proc. of the national academy of sciences 99, 12 (2002), 7821–7826.

[17] Ha, M., Baladandayuthapani, V., and Do, K. Dingo: differential network analysis in genomics. Bioinformatics 31, 21 (2015), 3413–20.

[18] Hamming, R. The unreasonable effectiveness of mathematics. American Mathematical Monthly 87, 2 (1980), 81–90.

[19] Horvath, S., Zhang, Y., Langfelder, P., Kahn, R. S., Boks, M. P., van Eijk, K., van den Berg, L. H., and Ophoff, R. A. Aging effects on DNA methylation modules in human brain and blood tissue. Genome biology 13, 10 (2012), R97.

[20] Huang, D. W., Sherman, B. T., and Lempicki, R. A. Systematic and integrative analysis of large gene lists using david bioinformatics resources. Nature protocols 4, 1 (2009), 44–57.

[21] Hubert, L. J. Assignment methods in combinatorial data analysis. Marcel Dekker 1 (1987).

[22] Ideker, T., Ozier, O., Schwikowski, B., and Siegel, A. Discovery regulartory and signalling circuits in molecular interaction networks. Bioinformatics 18 (2002).

[23] Jiao, Y., Widschwdter, M., and Teschendorff, A. E. A systems-level integrative framework for genome-wide dna methylation and gene expression data identifies differential gene expression modules under epigenetic control. Bioinformatics 30, 16 (2014), 2360–2366.

[24] Jin, L., Chen, Y., Wang, T., Hui, P., and Vasilakos, A. Understanding user behavior in online social networks: a survey. Communications Magazine, IEEE 51, 9 (September 2013), 144–150.

[25] Johnson, W. E., Li, C., and Rabinovic, A. Adjusting batch effects in microarray expression data using empirical bayes methods. Biostatistics 8, 1 (2007), 118–127.

[26] Jolma, A., Yan, J., Whitington, T., Toivonen, J., Nitta, K. R., Rastas, P., Morgunova, E., Enge, M., Taipale, M., Wei, G., et al. Dna-binding specificities of human transcription factors. Cell 152, 1 (2013), 327–339.

[27] Keller, A., Bakes, C., Gerasch, A., Kaufmann, M., Kohlbacher, O., Meese, E., and Lenhof, H. A novel algorithm for detecting differentially regulated paths based on gene enrichment analysis. Bioinfomatics 25, 21 (2009), 2787–2794.

[28] Kulakovskiy, I. V., Vorontsov, I. E.,Yevshin, I. S.,Soboleva, A. V.,Kasianov, A. S.,Ashoor, H.,Ba-Alawi, W.,Bajic, V. B.,Medvedeva, Y. A.,Kolpakov, F. A., et al. Hocomoco: expansion and enhancement of the collection of transcription factor binding sites models. Nucleic acidsresearch _44,_ D1 (2016), D116–D125.

[29] Lamirel, J.-C., Cuxac, P., Mall, R., and Safi, G. A new efficient and unbiased approach for clustering quality evaluation. New Frontiers in Applied Data Mining (2012), 209–220.

[30] Lena, P. D., Wu, G., Martelli, P., Casadio, R., and Nardini, M. C. An efficient tool for molecular interaction maps overlap. BMC Bioinforma 14, 1 (2013), 159.

[31] Levandowsky, M., and Winter, D. Distance between sets. Nature 234, 5323 (1971), 34–35.

[32] Li, D., Brown, J. B., Orsini, L., Pan, Z., Hu, G., and He, S. Moda: Module differential analysis for weighted gene co-expression network. arXiv preprint arXiv:1605.04739 (2016).

[33] Mall, R., Cerulo, L., Bensmail, H., Iavarone, A., and Ceccarelli, M. Detection of statistically significant network changes in complex biological networks. BMC Systems Biology 11, 1 (2017), 32.

[34] Mall, R., Langone, R., and Suykens, J. A. Kernel spectral clustering for big data networks. Entropy 15, 5 (2013), 1567–1586.

[35] Mall, R., Langone, R., and Suykens, J. A. Self-tuned kernel spectral clustering for large scale networks. In Big Data, 2013 IEEE International Conference on (2013), IEEE, pp. 385–393.

[36] Mall, R., Langone, R., and Suykens, J. A. Multilevel hierarchical kernel spectral clustering for real-life large scale complex networks. PloS one 9, 6 (2014),e99966.

[37] Mantel, N. The detection of disease clustering and a generalized regression approach. Cancer Research 27, 2 (1967), 209.

[38] Marbach, D., Lamparter, D., Quon, G., Kellis, M., Kutalik, Z., and Bergmann, S. Tissue-specific regulatory circuits reveal variable modular perturbations across complex diseases. Nature methods (2016).

[39] Margolin, A. A., Nemenman, I., Basso, K., Wiggins, C., Stolovitzky, G., Favera, R. D., and Califano, A. Aracne: An algorithm for the reconstruction of gene regulatory networks in a mammalian cellular context. BMC Bioinformatics 7, S–1 (2006).

[40] Mathelier, A., Fornes, O., Arenillas, D. J., Chen, C.-y., Denay, G., Lee, J., Shi, W., Shyr, C., Tan, G., Worsley-Hunt, R., et al. Jaspar 2016: a major expansion and update of the open-access database of transcription factor binding profiles. Nucleic acids research 44, D1 (2016), D110–D115.

[41] Merico, D., Isserlin, R., Stueker, O., Emili, A., and Bader, G. D. Enrichment map: a network-based method for gene-set enrichment visualization and interpretation. PloS one 5, 11 (2010), e13984.

[42] Mislove, A., Marcon, M., Gummadi, K. P., Druschel, P., and Bhattacharjee, B. Measurement and analysis of online social networks. In Proc. of the 7th ACM SIGCOMM Conference on Internet Measurement (2007), IMC ‘07, ACM, pp. 29–42.

[43] Nacu, S., Critchley-Throne, R., Lee, R., and Holmes, S. Gene expression network analysis and applications to immunology. Bioinformatics 23, 7 (2007), 850–858.

[44] Orman, G. K., and Labatut, V. A comparison of community detection algorithms on artificial networks. In International Conference on Discovery Science (2009), Springer, pp. 242–256.

[45] Pržulj, N. Biological network comparison using graphlet degree distribution. Bioinformatics 23, 2 (2007), e177–e183.

[46] Ramana, M., Scheinerman, E., and Ullman, D. Fractional isomorphism of graphs. Discrete Mathematics 132, 1 (1994), 247–265.

[47] Reichardt, J., and Bornholdt, S. Statistical mechanics of community detection. Physical Review E _74_, 1 (2006), 016110.

[48] Rosvall, M., and Bergstrom, C. T. Multilevel compression of random walks on networks reveals hierarchical organization in large integrated systems. PloS one 6, 4 (2011), e18209.

[49] Ruan, D. Statistical methods for comparing labelled graphs. PhD thesis, Imperial College London, 2014.

[50] Ruan, D., Young, A., and Montana, G. Differential analysis of biological networks. BMC bioinformatics 16, 1 (2015), 327.

[51] Shervashidze, N., Schweitzer, P., van Leeuwen, E. J., Mehlhorn, K., and Borgwardt, K. Weisfeiler-Lehman Graph Kernels. Journal of Machine Learning Research 12 (2011), 2539–2561.

[52] Teschendorff, A. E., Menon, U., Gentry-Maharaj, A., Ramus, S. J., Weisenberger, D. J., Shen, H., Campan, M., Noushmehr, H., Bell, C. G., Maxwell, A. P., et al. Age-dependent DNA methylation of genes that are suppressed in stem cells is a hallmark of cancer. Genome research 20, 4 (2010), 440–446.

[53] Wallace, T., Martin, D., and Ambs, S. Interaction among genes, tumor biology and the environment in cancer health disparities: examining the evidence on a national and global scale. Carcinogenesis 32, 8 (2011), 1107 1121.

[54] West, J., Beck, S., Wang, X., and Teschendorff, A. E. An integrative network algorithm identifies age-associated differential methylation interactome hotspots targeting stem-cell differentiation pathways. Scientific reports 3 (2013), 1630.

[55] Yang, Q., and Sze, S. Path matching and graph matching in biological networks. Journal of Computational Biology 14, 1 (2007), 56–67.

[56] Yang, X., Shao, X., Gao, L., and Zhang, S. Systematic dna methylation analysis of multiple cell lines reveals common and specific patterns within and across tissues of origin. Human molecular genetics 24, 15 (2015), 4374–4384.

[57] Zhang, B., Horvath, S., et al. A general framework for weighted gene co-expression network analysis. Statistical applications in genetics and molecular biology 4, 1 (2005), 1128.

